# Multivariate Residualization in Medical Imaging Analysis

**DOI:** 10.1101/2023.02.15.528657

**Authors:** Kevin Donovan, Nicholas J Tustison, Kristin A. Linn, Russell T. Shinohara, the Alzheimer’s Disease Neuroimaging Initiative

## Abstract

Nuisance variables in medical imaging research are common, complicating association and prediction studies based on image data. Medical image data are typically high dimensional, often consisting of many highly correlated features. As a result, computationally efficient and robust methods to address nuisance variables are difficult to implement. By-region univariate residualization is commonly used to remove the influence of nuisance variables, as are various extensions. However, these methods neglect multivariate properties and may fail to fully remove influence related to the joint distribution of these regions. Some methods, such as functional regression and others, do consider multivariate properties when controlling for nuisance variables. However, the utility of these methods is limited for data with many image regions due to computational and model complexity. We develop a multivariate residualization method to estimate the association between the image and nuisance variable using a machine learning algorithm and then compute the orthogonal projection of each subject’s image data onto this space. We illustrate this method’s performance in a set of simulation studies and apply it to data from the Alzheimer’s Disease Neuroimaging Initiative (ADNI).

## 1 Introduction

The presence of nuisance variables is a frequent complication in statistical analysis. Given variables *X* and *Y*, a nuisance variable *A* is one related to both *X* and *Y*. When not accounted for, nuisance variables can lead to erroneous conclusions about the relationship between two or more variables of interest. For example, influence of A may make *X* and *Y* to appear to be related, even if this relationship is entirely attributable to A. Furthermore, predictive models for *Y* based on *X* may not be generalizable across values of *A*, leading to incorrect conclusions about the predictive utility of *X* in the population [9, 7]. Thus, properly accounting for nuisance variables is critical in analyzing the relationships of interest.

Nuisance variables are common in medical imaging analyses, specifically when trying to analyze the predictive relationships between features of an image and an outcome variable (e.g., diagnosis) [9]. Imaging data are often high-dimensional, and features of the images are typically highly correlated. These correlations (along with location-specific means and variances) may differ depending on values of certain nuisance variables, which can induce spurious relationships between the image and outcome of interest [2]. We refer to this phenomenon as *multivariate confounding.* As a result, properly accounting for these nuisance variables may require taking into account the joint distribution of the image features, in addition to univariate information from each image feature.

Methodology for accounting for nuisance variables has been developed in the causal inference, predictive modeling, and machine learning literature. Methods accounting for nuisance variables generally fall into two categories: 1) those that adjust for nuisance variables and conduct analysis of interest in a single model (e.g deep learning or support vector machine) [15, 18, 17, 27] or 2) those that remove influence of the nuisance variables by transforming the predictors and use the transformed variables with the analytic method of choice [18, 19, 26, 2]. The most common of these is *univariate residualization,* which involves fitting a series of univariate linear models with the nuisance variables as the covariates and each image feature as the outcome. The residuals from these models are then used as adjusted images which are marginally uncorrelated with the nuisance variables [7]. This approach also retains the original dimension of the imaging data.

Methods that account for the joint distribution of the images generally fall into Category 1, and are thus tied to a specific downstream prediction model or method to reduce the dimension of the imaging data using a feature extraction approach such as principal components analysis (PCA) or partial least squares (PLS) [16, 26]. It would be useful to develop an adjustment method that is analogous to univariate residualization and falls into Category 2 which accounts for the potential multivariate confounding present in imaging data. A computationally efficient implementation of this adjustment would allow the use of adjusted images without being tied to a specific prediction analysis model, especially given the computational cost of some models (e.g. deep learning and functional regression). To our knowledge, such a method does not exist in the literature.

In this work, we develop a method to generate a set of adjusted predictors (e.g. images for example) that minimize the predictive relationship between the original predictors and nuisance variable at the univariate and multivariate level. This process is referred to as *multivariate residualization.* At the same time, the relationship between the residualized predictors and outcomes, which is independent of the nuisance variable, maintains as much predictive value as possible. The original dimension of the predictor space is also maintained and the method is computationally efficient for high-dimensional data.

One example of a nuisance variable in medical imaging is age in the context of studies of Alzheimer’s disease (AD). One marker of AD is atrophy in an individual’s cerebral cortex, which can be measured using structural magnetic resonance imaging (MRI). However, age has been shown to be associated with both AD prevalence and patterns of cortical thinning in healthy aging, making it an important variable to control for when modeling the association between brain structure and AD diagnosis [8, 14, 5, 9, 13]. Motivated by this challenge, the proposed method’s performance is also evaluated with an analysis of data from the Alzheimer’s Disease Neuroimaging Initiative (ADNI). Its performance is also evaluated in simulation studies, with univariate residualization as the primary comparator method.

In Section 2, we outline the proposed multivariate residualization methods and illustrate multivariate confounding. In Section 3, we present simulation studies which evaluate the performance of multivariate residualization and compare it to univariate residualization. In Section 4, we describe the ADNI dataset, the degree of age-related confounding in these data, and apply the proposed method. We provide discussion and conclusions in Section 5. Code for implementing the proposed multivariate residualization methods and all results in this manuscript is provided at https://github.com/kmdono02/Multivariate_Residualization.

## 2 Methods

### 2.1 Motivation

Let *X* and *Y* denote two related variables of interest and *A* denote a nuisance variable between *X* and *Y*. Let *X* be a multivariate vector of dimension *p* and let *Y* and A be univariate. It is assumed that these variables have some degree of interdependence, visualized in the following diagram, with double-sided arrows denoting covariation between the variables conditional on the other, with these conditional covariances denoted by *γ_XY_, γ_AY_*, and *γ_AX_*.

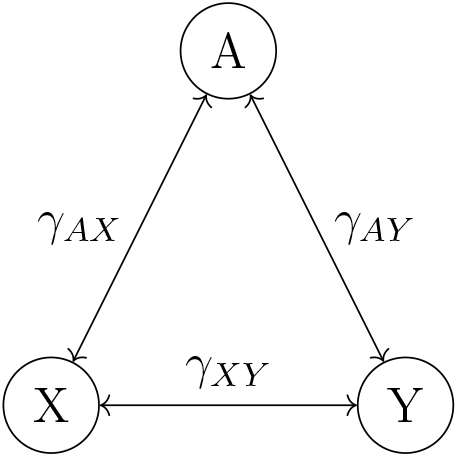

As an example, *A* may denote a person’s age, *X* may denote measurements from a set of medical imaging regions of interest (ROI), and *Y* may denote a disorder or disease that is associated with *X*. *γ_XY_* is considered of interest as it represents the covariation between the predictors and outcome independent of their relationships with the nuisance variable. The goal is to remove the influence of *γ_AX_* from the images while maintaining *γ_XY_*.

Depending on the data-generating mechanism corresponding to this diagram, traditional methods such as univariate residualization can successfully remove the influence of *A*. Let {*x_i_,a_i_,y_i_*} for *i* =1, …,*n* denote the realized predictor vectors, nusiance variables, and outcomes for a random sample of size n. Suppose *a_i_* ∈ (-∞, ∞) and *y_i_* ∈ {0,1} ∀*i*. To consider univariate residualization in the context of a downstream prediction analysis for *Y*, we define univariate residualization in the context of a separate training and testing set. Let indices *i* = 1,…, *n_t_* denote the training set and *i* = *n_t_* +1, …, *n* denote the testing set. First, we define univariate per-predictor linear models on the training set by *x_ij_* = *β*_0*j*_ + *β*_1*j*_ *a_i_* + *ϵ_ij_* for *i* = 1, …, *n_t_* and *j* = 1, …, *p*. These models are fit using the training set only, with the estimated coefficients denoted by vectors 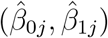, for *j* = 1, …, *p*. Then residuals for the whole sample are defined by 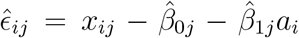 for *i* = 1, …, *n* and *j* = 1,…, *p*. The output of univariate residualization is defined here by p-dimension vectors 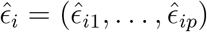 for *i* = 1, …, *n*.

If the joint distribution of *X* is independent of *A* and the variance of each element of *X* is independent of *A*, then by-location linear models can be used successfully. This is because the residuals will be uncorrelated with the outcome variable in a regression model by design of linear model fitting, and the model’s assumption of homoscedasticity will be satisfied. However, if the association with *A* is also evident in the joint distribution of *X*, for example in the correlation between image locations, then this association will not be removed by univariate residualization. As a result, this remaining association can be detected and used by prediction algorithms, biasing the estimated relationship between *X* and *Y* [6]. This can be seen in the following simulated example shown in Figure 1; the correlation between *X*_1_ and *X*_2_ differs if *A* is below 0 or above 0 [6]. Thus, *A* is related to the joint distribution of *X*_1_ and *X*_2_, despite *X*_1_ and *X*_2_ being uncorrelated marginally with the same means and variances. See Section 3.1 for details on this simulation.

**Figure 1:**
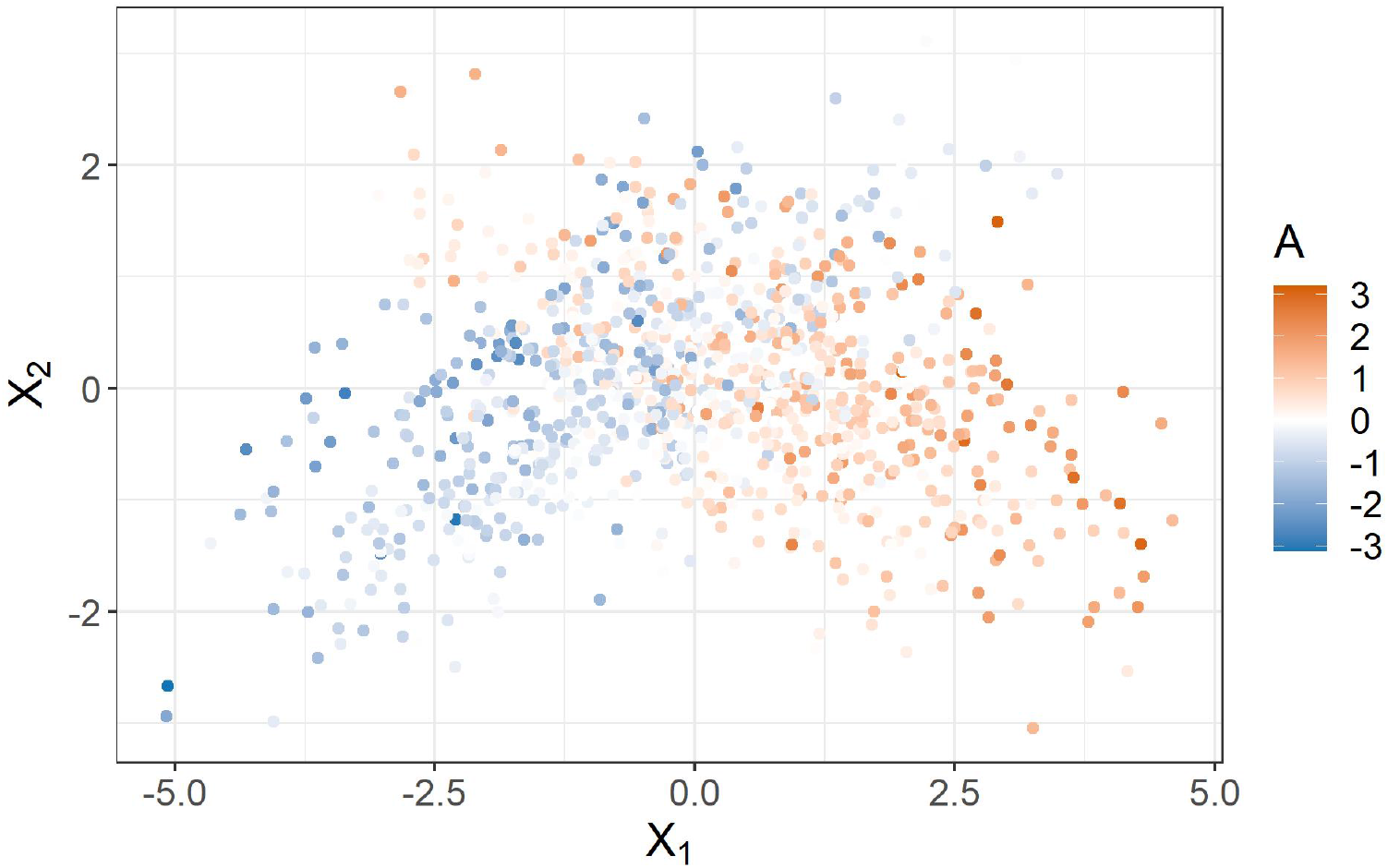
Multivariate confounding simulated example. Correlation between *X*_1_ and *X*_2_ differs based on *A*, though means and variances of *X*_1_ and *X*_2_ are equal.

### 2.2 Multivariate Residualization

The first step of the proposed method is to estimate the predictive relationship between *A* and *X* jointly, e.g. the covariation between the multivariate image *X* and *A*. We propose modeling this relationship with a linear support vector regression (SVR) model with *A* as the outcome and *X* as the predictor vector. SVR is used due to its ability to handle *p* larger then *n*, flexibility in handling outliers using the tuning parameters, and computational efficiency and interpretability [24].

A linear SVR is defined as follows. Let *i* = 1,…, *n_t_* denote the indices for a training subset of size *n_t_* from a sample of size n. Suppose {*x_i_,α_i_,y_i_*} for *i* = 1, …, *n* are again observed in the sample with the goal of predicting *A* using *X* in a linear prediction model. For each *x_i_*, define the *i^th^* observation’s predicted *A* by 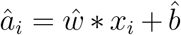 where *ŵ* denotes the estimated “weights” and 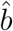 denotes the estimated intercept with * denoting a dot product. Let *ϵ*^+^ and *ϵ*^−^, both non-negative, denote slack variables with margin *ϵ* > 0. To compute *ŵ* and 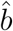, the following optimization problem is solved using the training set’s observations:

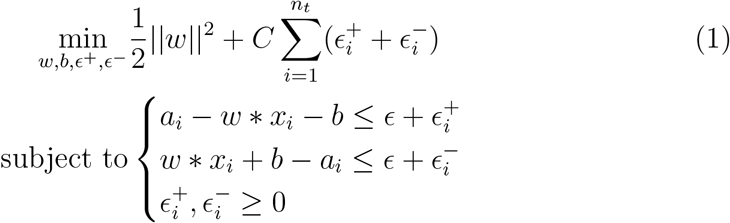

where *C* and *ϵ* > 0 are hyperparameters to be chosen. Estimates *ŵ* and 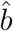 are the values of *w* and *b* respectively which solve Equation 1. See [24] for details.

One potential concern is that *Y* is not accounted for in this optimization routine to obtain *ŵ*. As a result, the estimated relationship between *X* and *A* may be biased (i.e., Simpson’s Paradox [1]). Thus, we propose to fit separate SVR models in each *Y* group. Assume *Y* is binary for simplicity (though this process could be implemented with multinomial *Y*). Let *ŵ*_0_ and *ŵ*_1_ denote the SVR weights obtained by solving the optimization problem 1 for *w*, using only observations in the training data where *Y* = 0 and *Y* = 1 respectively.

After solving for *ŵ*_0_ and *ŵ*_1_, we compute the orthogonal projection of each observation’s *x_i_* vector to the mean of these *Y*-specific weight vectors. This step is analogous to extracting linear regression residuals which are orthogonal to the fitted values and thus uncorrelated with the model’s covariates [12]. Let 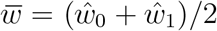, a vector of length *p* denoting the average of the SVR weight vectors from the *Y* groups. Then, for a subject’s observed *x_i_*, define 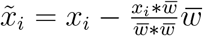 for *i* = 1,…, *n*. This is the orthognal projection of *x_i_* onto 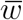, which we denote as this subject’s residual in the context of multivariate residualization to *A*. Thus, new observations can be orthogonalized without knowing their observed *Y*. This process is summarized in Algorithm 1, labeled *SVR-MR*. The hyperparameters for the SVR are tuned in the training set fits, using a grid search or similar approach to minimize mean squared error (MSE) between *X* and *A* with K-fold cross validation. This results in a set of residuals 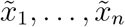, each of dimension *p*, which aim to satisfy 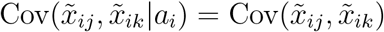 for *j* = *k, j, k* = 1,…, *p* and *i* = 1,…, *n*.

#### Algorithm 1: SVR-based multivariate residualization (SVR-MR)

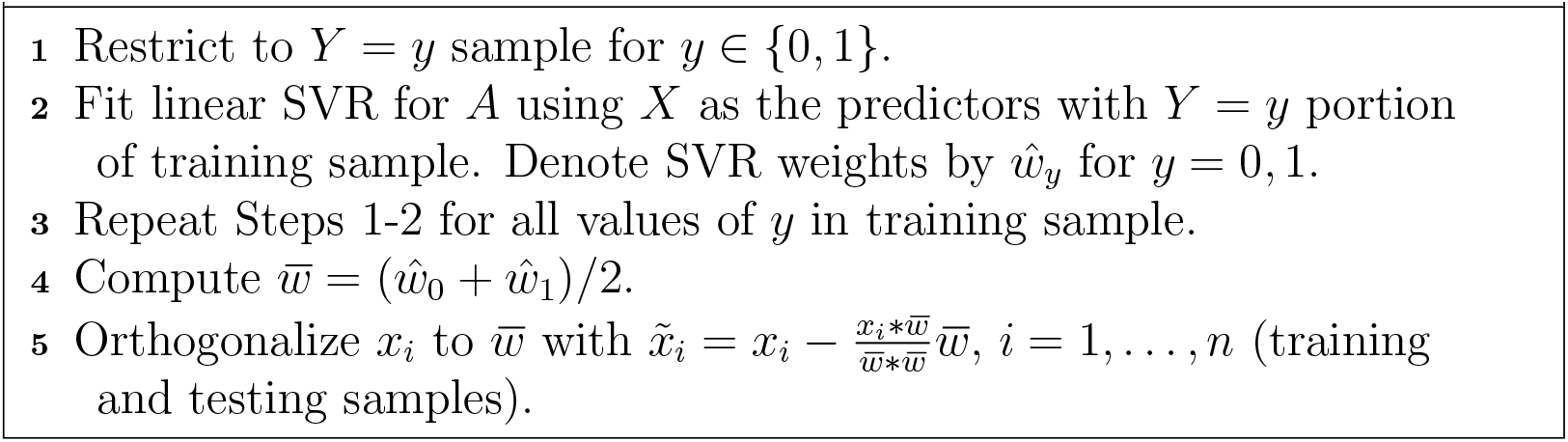

### 2.3 Extensions

While the residuals from SVR-MR are designed to have associations between *X* and *A* removed at the covariance level, marginal associations between *X* and A may still remain. This is caused by the use of all *X* jointly in the SVR fit step. Since all variables in multidimensional *X* are controlled for when fitting the SVR model, some predictors’ SVR weights may be near zero even though these predictors are correlated with *A*. This can occur if, for example, *X*_1_ is marginally correlated with *A*, however uncorrelated with *A* conditional on the other *X*. As a result, such predictors are minimally adjusted after SVR-MR, resulting in remaining marginal correlations with *A*. We develop extensions of SVR-MR to additionally remove these remaining correlations. An illustrative example is shown in Section 3.2.

The first extension, labeled *SVR Repeat,* applies SVR-MR multiple times until the change in training set *R*^2^ when predicting *A* with the latest round’s residuals is smaller then a chosen threshold (5% change was used). That is, each previous round’s residuals are residualized again using the same SVR-MR process, until the threshold is reached. The second extension, labeled *SVR + UR* denotes one round of SVR-MR followed by univariate residualization with the SVR-MR residuals as the outcomes in the per-residual univariate residualization models. These new residuals are extracted as the final residualized images for each subject. These extensions result in a set of residuals, again labeled 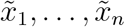, each of dimension *p*, which aim to satisfy both the condition for SVR-MR in 2.2 and 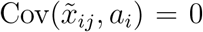 for *j* = *k,j,k* = 1,…, *p* and *i* = 1,…, *n*.

## 3 Simulation Studies

In this section, we examine the proposed residualization methods using simulation studies. In each study, we compare the methods’ performance to univariate residualization in terms of predicting *A* and *Y* after residualization. Two settings for the distributions of these simulated data are considered: 1) Cov(*X_j_, X_k_* |*A*) differs for values of *A* (multivariate confounding) and 2) Cov(*X_j_,X_k_*|*A*) is constant across *A*. In both settings, *Y* is independent of *X* conditional on *A*. The residualization methods are then evaluated in these simulations based on the associations between their corresponding residuals and *A* as well as with *Y*. Ideally, these residuals should have minimal predictive associations with the nuisance variable *A*, with a higher MSE indicating improved residualization performance. In terms of binary *Y*, if *X* and *Y* are independent conditional on *A*, these residuals should fail at predicting *Y*. That is, an area under the curve (AUC) for a corresponding prediction model around 0.5 would be expected.

The structure for these studies is as follows, visualized in Figure 2. In all simulations, *n* = 1000 with the following analysis pipeline. First, we randomly partition 50:50 the simulated sample into two sets balanced by *Y*, denoted training and evaluation sets respectively. The training set is used to train the residualization methods and the evaluation set is used to quantify the performance of these methods. We trained the SVR-MR model with the training set’s data partitioned by *Y* with *A* as the outcome and all *X* as the predictors, using the process from Section 2.2. When training SVR Repeat, we compute residuals in the training set using one round of SVR- MR, and then the SVR-MR process was repeated using these residuals as the predictors, again partitioned by *Y*. This process ends once the *R*^2^ computed from these residuals and *A* in the training set was below 5%, per Section 2.3. This *R*^2^ is computed after each round based on predictions from a linear SVR with *A* as the outcome and the round’s residuals as predictors. We train univariate residualization and SVR+UR in the same fashion. For each method, we extract the final weights from the fitting process to use with the evaluation set.

**Figure 2:**
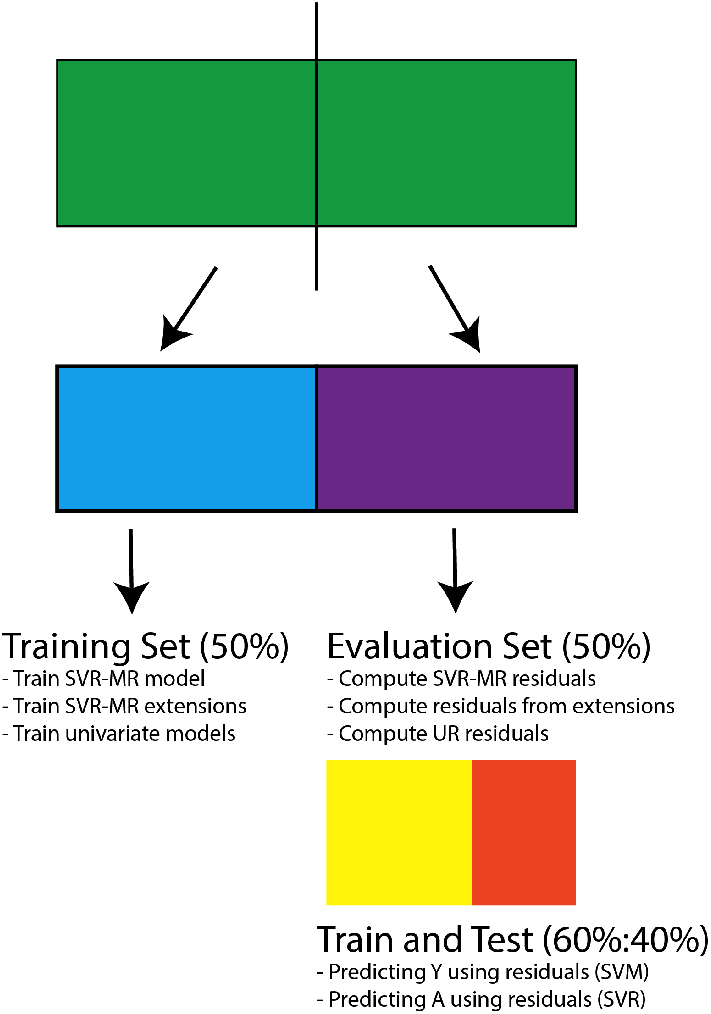
Diagram of data splitting and analysis pipeline used for simulation studies.

With the evaluation set, for each method, we compute residuals using the final weights (SVR weights and/or regression coefficients) from the above process. This is used to examine the out-of-sample generalizability of these residualization methods to samples which are not used to estimate the models for residualization. After computing the residuals using these weights with the process from Section 2.2, we compare the residualization methods in terms of predicting *A* and predicting *Y*. In the evaluation set, we generate training and testing sets using a random 60:40 split of the data balanced by *Y*. To assess the residuals’ predictive association with *A*, a radial SVR model is tuned and fit to the training set specified above with *A* as the outcome and the residuals as the predictors. We compute the out-of-sample MSE for *A* with the testing set as a metric of predictive association. To assess the residuals’ predictive association with *Y*, a radial support vector machine (SVM) model is tuned and fit to the training set specified above with *Y* as the outcome and the residuals as the predictors. This is repeated for each method’s set of residuals. We compute the out-of-sample AUC for *Y* with the testing set as a metric of predictive association.

We repeat this process 1000 times for each simulation scenario. The out-of-sample MSE and AUC distributions across the different iterations are then compared across the different residualization methods, along with using *X* without adjustment to predict *A* and *Y* using the same training-testing scheme on the evaluation set as specified above.

### 3.1 Multivariate Confounding Setting

For the fist set of simulations, we consider the example shown in Figure 1. The data-generating mechanism for these simulations is the following:

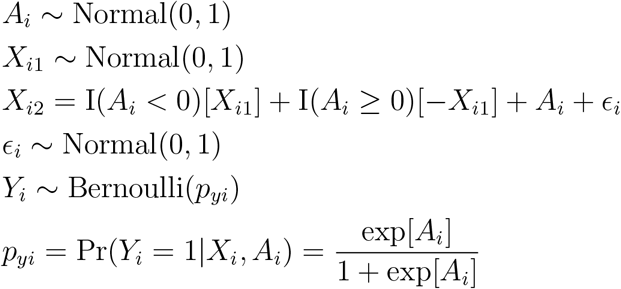

for *i* = 1,…, *n*, where *I*(·) denotes the indicator function. This data generating mechanism corresponds to *X* being independent of *Y* conditional on *A* (but not marginally independent of *Y*). That is, ideally a set of residualized *X* vectors would have no predictive relationship with *Y*.

Boxplots of the out-of-sample MSEs and AUCs across the 1000 repetitions are provided in Figure 3. Top row in each subfigure corresponds to using the unadjusted *X* to predict *A* and *Y*, followed by SVR-MR, repeated SVR-MR (SVR Repeat), SVR-MR then univariate residualization (SVR+UR), and lastly univariate residualization (UR). Using the unadjusted *X* results in a median AUC elevated from 0.5, with univariate residualization showing a median AUC of about 0.58. The methods that use SVR-MR show a distribution of AUC centered closer to 0.5, the accurate null value given that *X* was independent of *Y* controlling for *A* in these simulations. These results are also reflected in the MSE for *A*; the unadjusted *X* and univariate residualization show the lowest MSE while the multivariate residualization methods show elevated MSE with all showing similar distributions.

**Figure 3:**
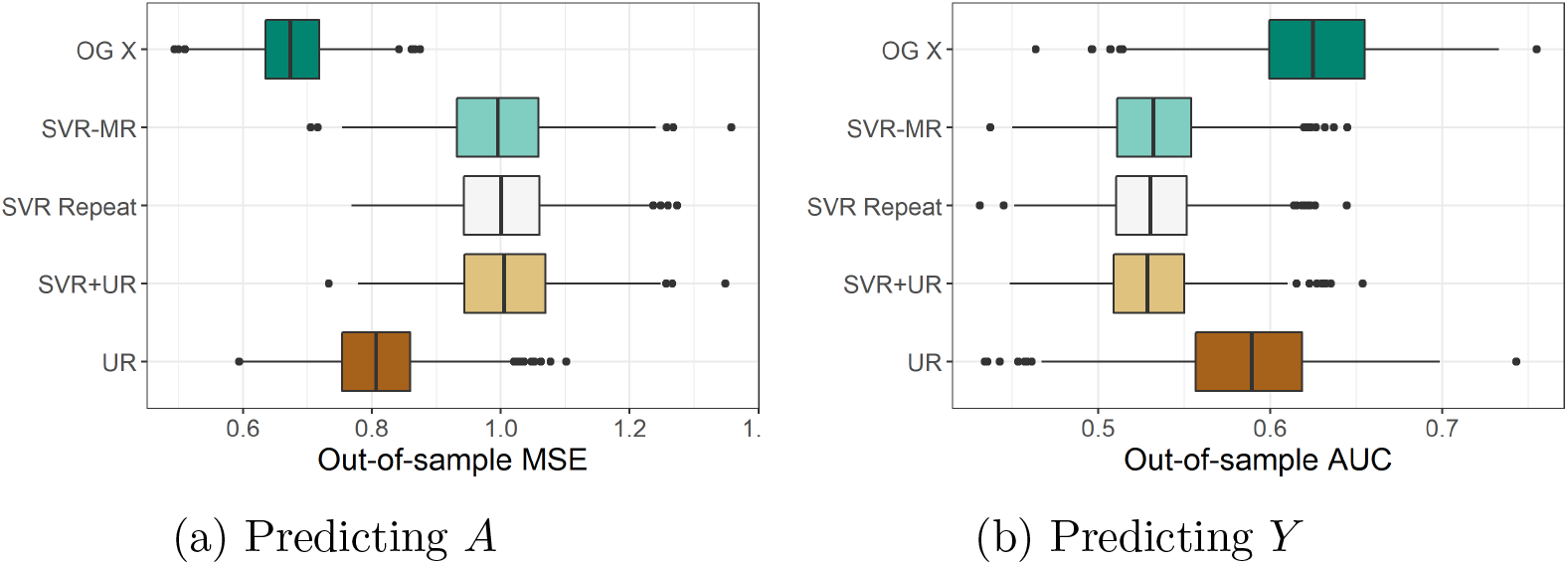
Distributions of out-of-sample MSE for predicting *A* (left) and AUC for predicting *Y* (right) using various residualization methods with the multivariate confounding scenario and 1000 repetitions. *OG X* denotes using unadjusted *X*. UR denotes univariate residualization.

### 3.2 Correlated Predictors Setting

Next we consider examples where *p* varies and correlations between *X* and *A* are solely through marginal associations between the predictors and nuisance variables. For these simulations, the data generating mechanism is the following:

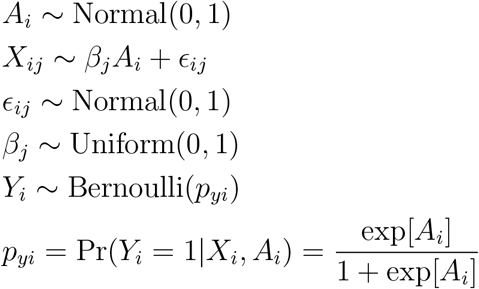

for *i* = 1, …, *n* and *j* = 1, …, *p*, with *p* = 5 and *p* = 50 considered. Slope parameters *β*_1_, …, *β_p_* are fixed after *p* independent draws for all simulations. This induces varied correlations between different elements of *X* depending on the magnitude of the slope parameters. In this context, a high MSE for predicting *A* and AUC near 0.5 would again be desired after any residualization. Since no multivariate confounding exists in these simulations, univariate residualization should do well.

Boxplots of the out-of-sample MSEs and AUCs across the 1000 repetitions for the different levels of *p* are provided in Figure 4. As expected, univariate residualization performs well in both contexts, with high MSE relative to using the unadjusted *X* and AUC centered close to 0.5. For *p* = 5, all multivariate residualization methods perform similarly to each other and similarly to univariate residualization. For *p* = 50, one round of SVR-MR performs pooly, with a lower increase in out-of-sample MSE relative to the other residualization methods and an AUC centered away from 0.5. However, SVR Repeat (with a 5% *R*^2^ threshold) and SVR+UR perform well and very similarly to univariate residualization. This is due to high marginal correlations with *A* remaining after SVR-MR due to the high correlations between the different *X* variables. Univariate residualization removes these marginal correlations since it does not control for all *X* when fitting the models, unlike a single round of SVR-MR.

**Figure 4:**
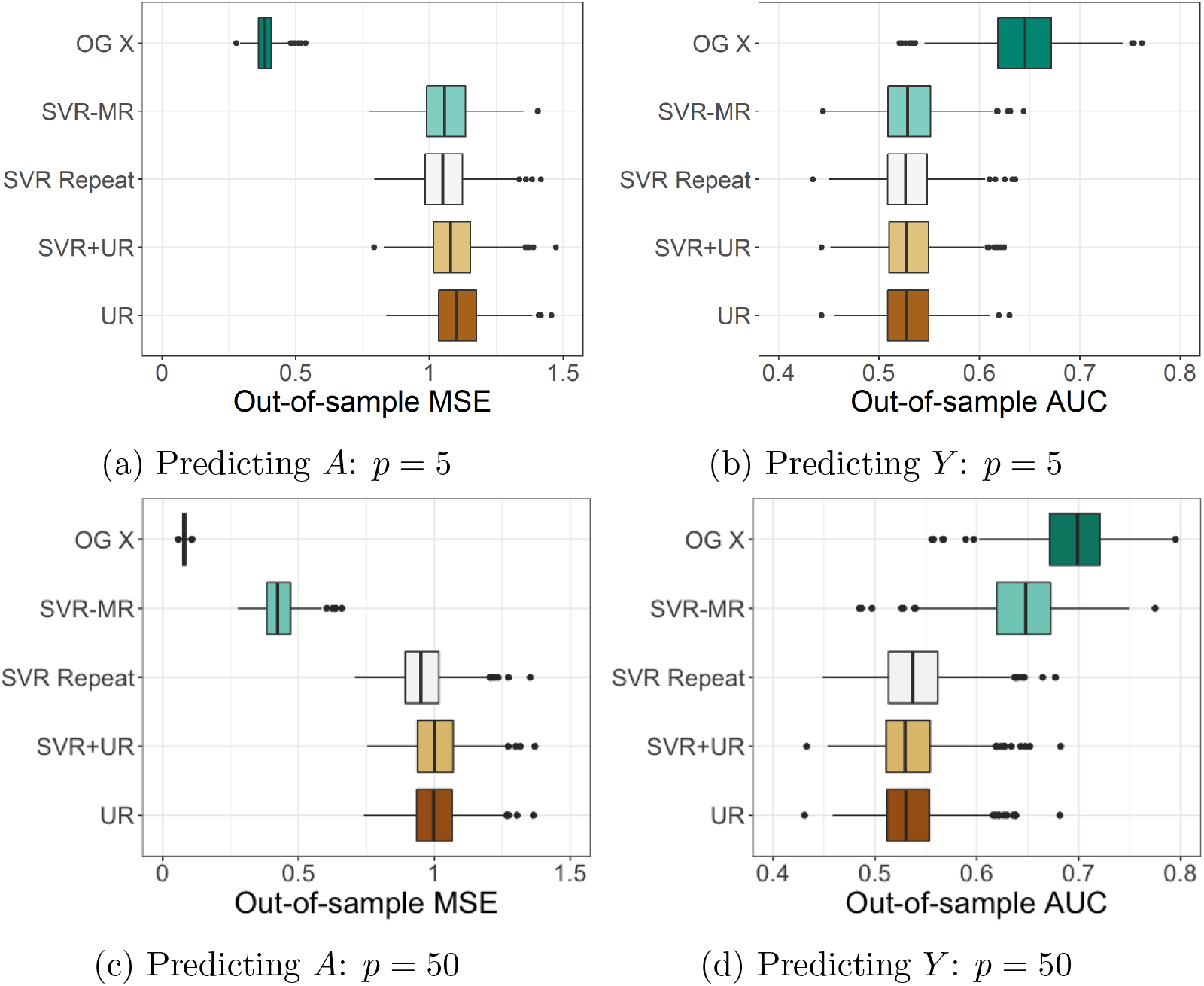
Distributions of out-of-sample MSE for predicting *A* (left) and AUC for predicting *Y* (right) using various residualization methods with the correlated predictors scenarios and 1000 repetitions. Top row of plots denotes *p* = 5 results, bottom *p* = 50 results. *OG X* denotes using unadjusted *X*. UR denotes univariate residualization.

## 4 ADNI Data Analysis

To evaluate the proposed methods in real data, we analyze data from the Alzheimer’s Disease Neuroimaging Initiative (ADNI) (https://adni.loni.usc.edu/). The ADNI was launched in 2003 as a public–private partnership, led by Principal Investigator Michael W. Weiner, MD. The main aim of ADNI has been to test whether longitudinal magnetic resonance imaging (MRI), positron emission tomography, other biomarkers, and clinical and neuropsychological assessment can be used together to accurately measure the progression of mild cognitive impairment (MCI) and early Alzheimer’s Disease (AD).

One biomarker for AD which has been established in the literature based on analysis of ADNI data is thickness of the cerebral cortex. After parcellation of the cortex into regions of interest (ROIs), multivariate analyses using average thickness in each ROI as predictors have shown predictive utility for AD and MCI diagnoses from healthy control (CN) [22, 10, 23]. However, reduction in cortical thickness is also associated with healthy aging in CN individuals. This may lead to biased conclusions from such multivariate analyses if age is not properly taken into account in the analysis [8, 14, 5, 9, 13]. Motivated by this work, we assess the residualization methods’ performance with the ADNI data in the context of predicting AD diagnosis with cortical thickness as predictors and age as a nuisance variable.

The data extracted for use in this analysis consist of structural MRI scans at each participant’s baseline visit, and is restricted to those with an AD diagnosis or considered a healthy control. From the baseline structural MRI scans, cortical thickness measurements from 60 ROIs for each participant are extracted [25]. This results in 122 AD and 163 CN participants with complete ROI cortical thickness data at baseline visit.

The ADNI data was age-matched, limiting the association between age and AD diagnosis present in the sample, in contrast to the association present in the general population. As a result, we create an age-biased subsample to induce an age-diagnosis association to assess the residualization methods with the ADNI data. This is done by randomly selecting 60 percent of the AD and CN individuals, where the probability of being selected was exp[4 · Age]/(1 + exp[4 · Age]) for AD observations and exp[−4 · Age]/(1 + exp[−4 · Age]) for CN observations. The differences in the age distributions between the AD and CN groups in the observed sample compared to the biased subsample are shown in Figure 5a, with the subsample showing elevated median age in AD compared to CN as is the case in the general population.

**Figure 5:**
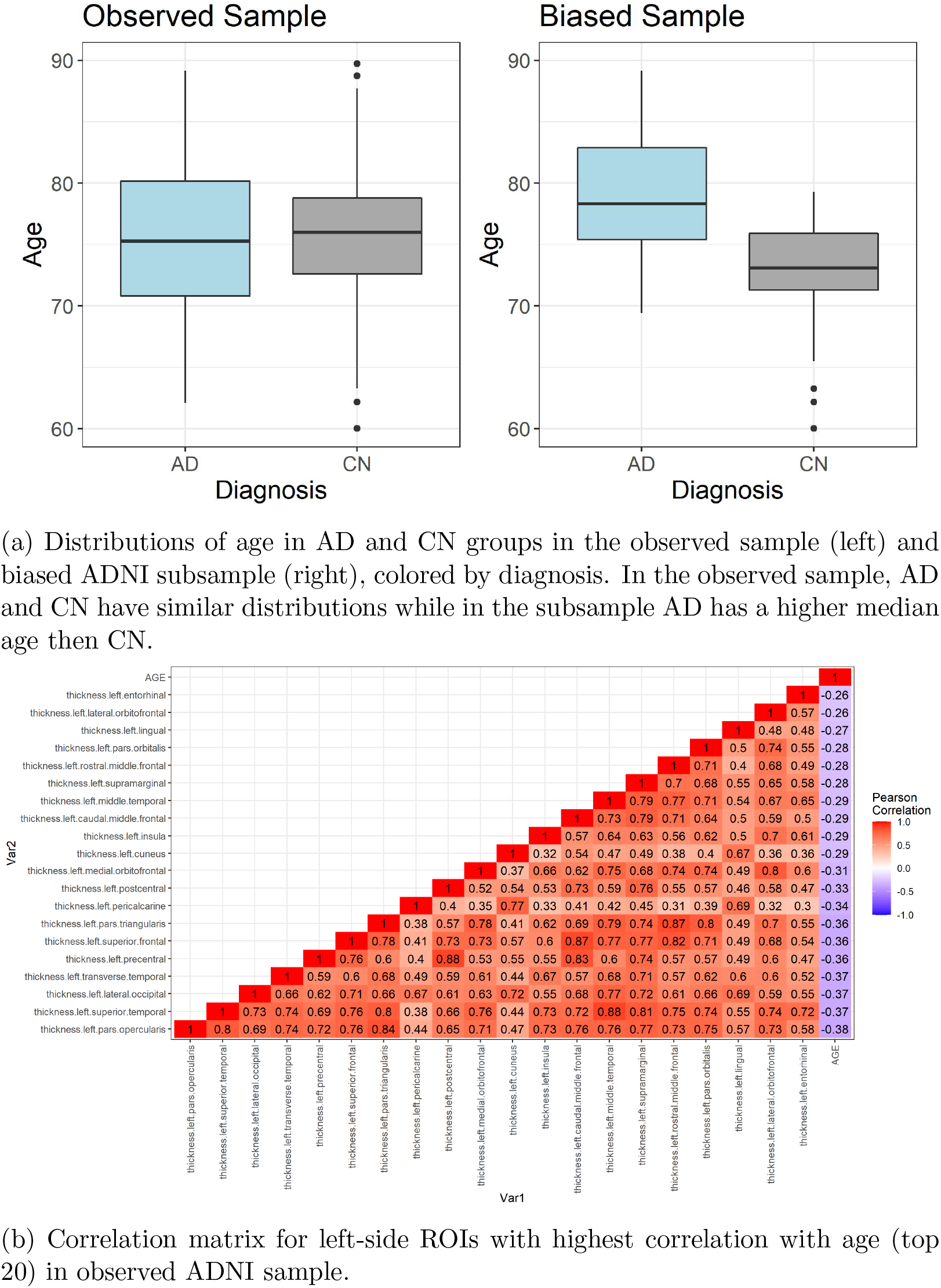
Summary statistics of observed ADNI data and biased ADNI data generated by subsampling. Distributions of age by diagnosis and cortical thickness correlations visualized.

The residualization methods examined in Section 2 are used to remove the influence of age on these ROIs in the biased subsample, and then assess the predictive utility of the ROIs for AD diagnosis after residualization. The 60 ROIs in the data consist of 30 locations, measured on the left and right side. Due to the high correlation between two hemispheres for each location, we only consider only one side’s ROIs (left). For ease of visualization, we use the 20 of these 30 ROIs with the highest absolute correlation with age as the final set of ROIs to evaluate the residualization methods. The estimated correlation matrix in the observed sample is provided in Figure 5b, showing that all 20 ROIs had a moderate negative correlation with age and high correlation amongst themselves.

We evaluate the performance of these methods in removing age-related confounding using out-of-sample MSE for predicting age and out-of-sample AUC for predicting diagnosis with the ROIs after residualization. These metrics are computed using the follow process. First, we randomly split the biased subsample 60:40 into the training and testing set respectively. Then, we fit the different residualization models on the training set to the 20 ROIs as described in Section 2, with residuals computed in the training and testing sets using these models fit on the training set only. Finally, a radial SVR and SVM are fit to the training data to predict age and diagnosis respectively using these residuals. We apply these prediction models to the testing set with out-of-sample MSE and AUC computed. We repeat this process 100 times with 100 random splits.

Figure 6 shows the distributions of these MSEs and AUCs across the 100 random splits as boxplots, with the individual split values super-imposed. Using one round and twice-repeated SVR-MR (SVR-MR and SVR Two respectively) show lower out-of-sample MSE and higher out-of-sample AUC relative to methods which incorporate univariate residualization. This is likely due to the high correlation between the ROIs, resulting in marginal associations remaining. However, all methods which use univariate residualization showed very similar MSE and AUC distributions. For all methods, the AUC distribution is centered away from the null value of 0.5, which is expected given the established relationship between cortical thickness and AD diagnosis. This similarity in performance between univariate residualization and SVR+UR may be a result of a lack of multivariate confounding being present in the sample, as in the simulation results in Section 3.2.

**Figure 6:**
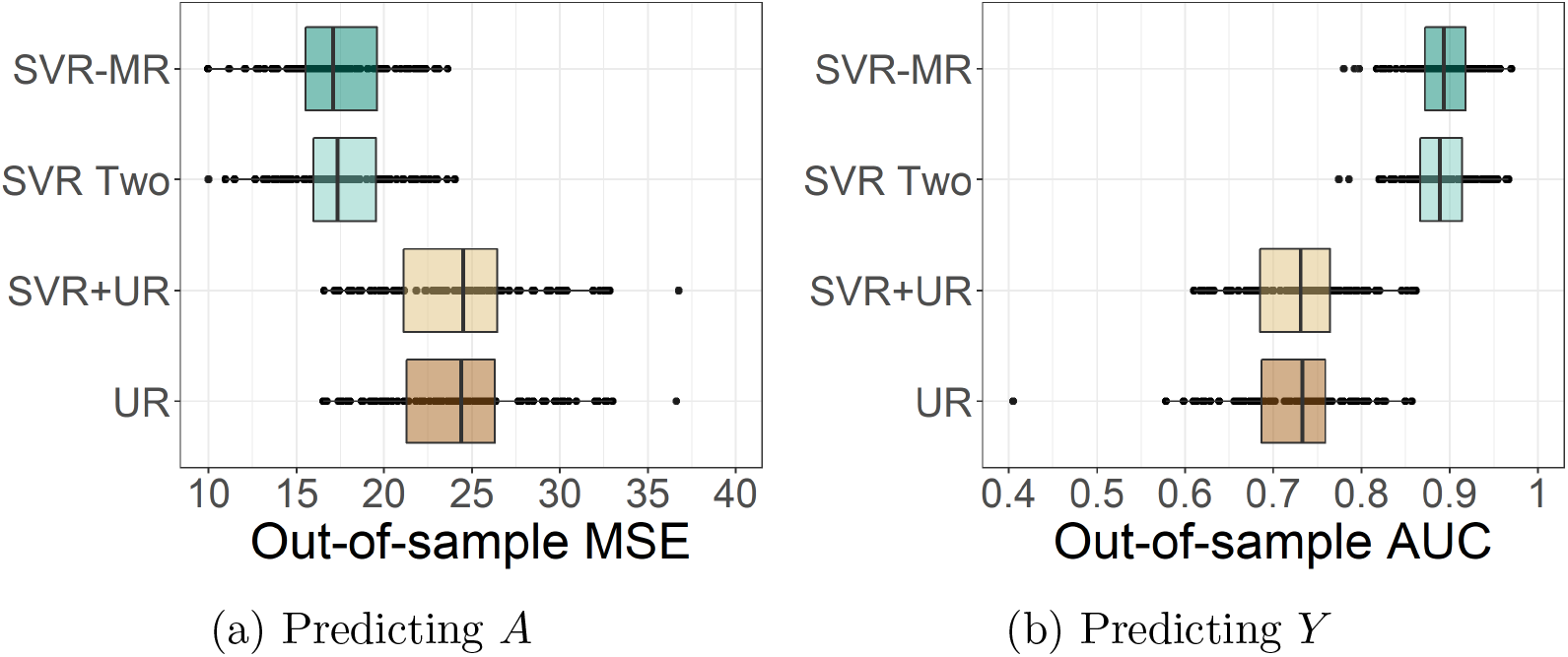
Distributions of out-of-sample MSE for predicting age (left) and AUC for predicting AD (right) using various residualization methods with the biased ADNI subsample and 100 repeated splits. UR denotes univariate residualization.

To investigate this similarity, we consider a set of simulation studies with the ADNI data in which multivariate confounding was induced. That is, let *X*_1_ denote the left-side ROI with the highest correlation with age in the biased subsample, normalized to mean 0 and variance 1. Let *Age* denote normalized age and *θ_Age_* denote median age after normalization. Then, we generate a second ROI for each observation in the biased subsample, *X*_2_, as

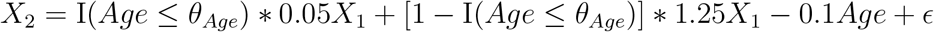

with *ϵ* generated from Normal(0,1). This creates a pair of ROIs that exhibit multivariate confounding with age, are positively correlated with each other marginally and conditionally on age, and negatively correlated with age, as was present in the original ROIs. These ROIs in the sample are visualized in Figure 7a. We conducted the same training and testing process that was implemented with the biased subsample using these two ROIs as the predictors in all fitted models and age again as the nuisance variable. We again repeated this process 100 times for 100 different randomly selected splits.

**Figure 7:**
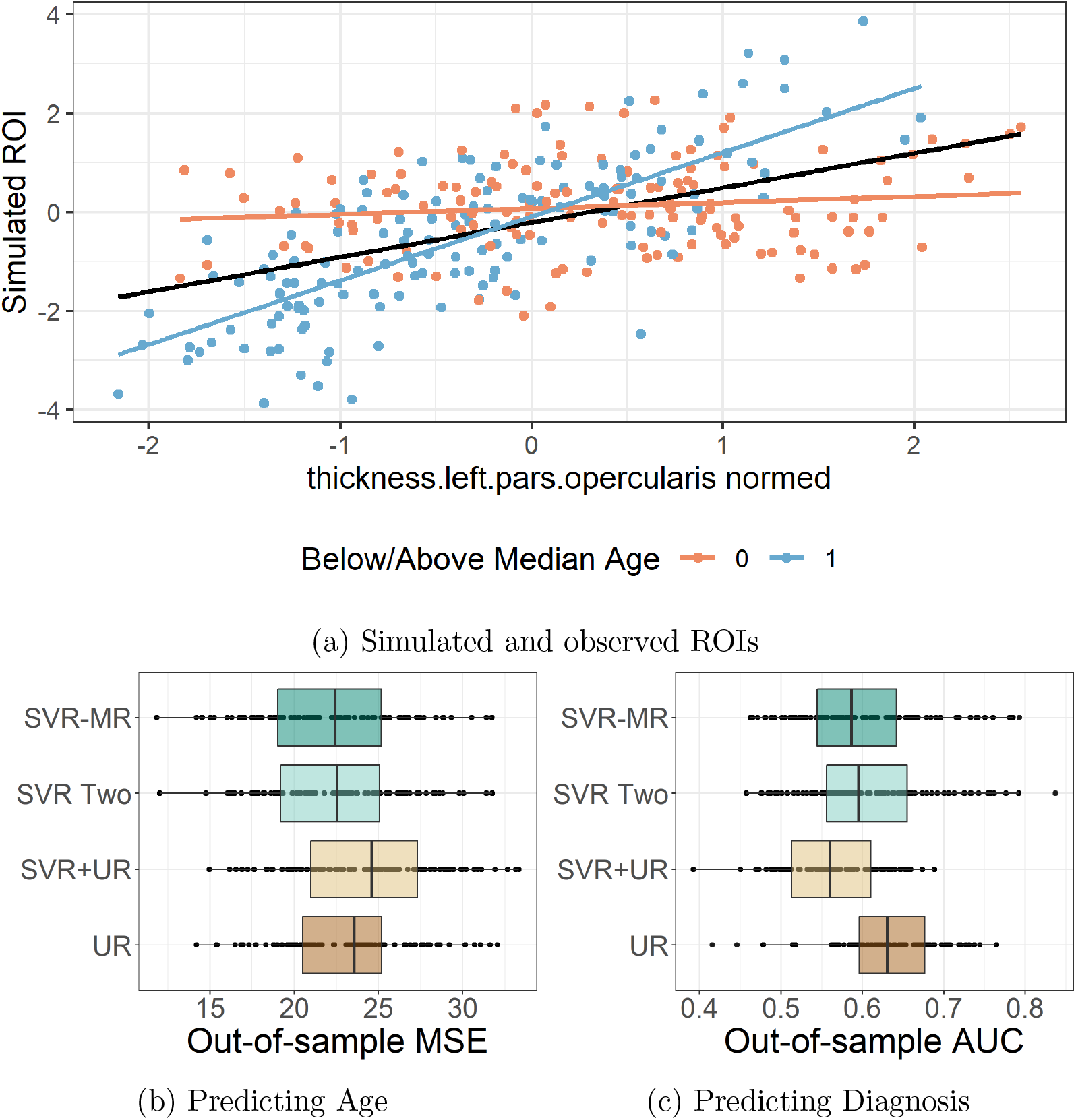
Results from analysis of biased ADNI subsample with simulated multivariate confounding. a) Scatterplot of observed and simulated ROIs in ADNI data, colored by age group (below or above median). Observed ROI was normalized to have mean 0 and variance 1. Lines of best fit for the two ROIs provided marginally (black line) and by age group. b) and c) Distributions of out-of-sample MSE for predicting age (left) and AUC for predicting AD (right) using various residualization methods with the biased ADNI subsample and 100 repeated splits. UR denotes univariate residualization.

Figures 7b-7c show the distributions of these MSEs and AUCs with the simulated ROI scenario. In this scenario, the two ROIs have non-zero marginal correlations with each other and differing conditional correlations with each other depending on age. For predicting age, one round or two rounds of SVR-MR and univariate residualization show similar out-of-sample MSEs when predicting age using the respective residuals. SVR+UR shows increased out-of-sample MSEs when predicting age when compared to using either type of method alone. Similar results are seen when predicting diagnosis with the residuals, where out-of-sample AUC decreased when using SVR+UR compared to using either type alone.

## 5 Discussion

In this paper, we develop a method to compute residuals which removes multivariate relationships between a set of predictors and a nuisance variable. This method is analogous to the commonly-used univariate residualization procedure, with the additional removal of nuisance variable associations with the covariance structure of the predictors, referred to here as multivariate confounding. Furthermore, like univariate residualization, the original dimension of the predictors is maintained and the method is computationally efficient for high dimensional predictors. While marginal correlations between the predictors and nuisance variable may remain for highly correlated predictors after SVR-MR, this can be remedied by performing univariate residualization on the residuals obtained from SVR-MR.

We evaluate these multivariate residualization methods and compare them to the univariate analogue using a set of simulation studies. These experiments show their ability to remove covariance relationships between the predictors and nuisance variable which the univariate method was unable to accomplish. In the case where only marginal association-related confounding was present, both univariate residualization and SVR+UR methods perform similarly, including when the predictors are correlated with one another. Finally, we use these methods to remove the association between age and cortical thickness ROIs in a sample from the ADNI database. Both univariate residualization and SVR+UR perform similarly when analyzing the observed ROIs. When simulating a ROI with multivariate confounding, SVR+UR shows improved performance in removing the association with age.

This method has several limitations. First, only one nuisance variable can be modeled at a time with the multivariate methods, compared to multiple nuisance variables with the univariate method. Also, each round of SVR-MR decreases the degrees of freedom for the set of residualized predictors by 1, effecting downstream analyses, particularly regression-based analyses with the residuals. This can be mitigated by using SVR+UR to remove remaining marginal associations between the SVR-MR residuals and the nuisance variable, instead of doing multiple rounds of SVR-MR.

Opportunities for future research include multivariate residualization simultaneously to multiple nuisance variables. While one could compute residuals using SVR-MR sequentially for each nuisance variable, this may not result in residuals which are jointly orthogonal to all of the nuisance variables. Another extension could be incorporating nonlinear relationships between the predictors and nuisance variables within the multivariate residualization. Nonlinear kernels within the SVR component of the proposed residualization method could considered for this purpose, though this would complicate the orthogonal projection step.

While univariate residualization and SVR+UR perform similarly when applied to ADNI data, the proposed multivariate residualization methods show promise for applications with AD and other neurological conditions. Previous research has found associations between the correlations within imaging features and diagnosis for various conditions, as well as nuisance variables such as age. Examples of these findings include both structural and functional connectivity and diagnosis, as well as age in AD and autism spectrum disorder [4, 20, 3, 11, 21]. The performance of our proposed methods in simulation studies indicate the potential utility of multivariate residualization when used in such examples. Investigating this in data from additional neurological studies would be useful to potentially better isolate associations between imaging features and diagnosis from nuisance variables.

## 7 Acknowledgements

Data collection and sharing for this project was funded by the Alzheimer’s Disease Neuroimaging Initiative (ADNI) (National Institutes of Health Grant U01 AG024904) and DOD ADNI (Department of Defense award number W81XWH-12-2-0012). ADNI is funded by the National Institute on Aging, the National Institute of Biomedical Imaging and Bioengineering, and through generous contributions from the following: AbbVie, Alzheimer’s Association; Alzheimer’s Drug Discovery Foundation; Araclon Biotech; BioClinica, Inc.; Biogen; Bristol-Myers Squibb Company; CereSpir, Inc.; Cogstate; Eisai Inc.; Elan Pharmaceuticals, Inc.; Eli Lilly and Company; EuroImmun; F. Hoffmann-La Roche Ltd and its affiliated company Genentech, Inc.; Fujirebio; GE Healthcare; IXICO Ltd.; Janssen Alzheimer Immunotherapy Research & Development, LLC.; Johnson & Johnson Pharmaceutical Research & Development LLC.; Lumosity; Lundbeck; Merck & Co., Inc.; Meso Scale Diagnostics, LLC.; NeuroRx Research; Neurotrack Technologies; Novartis Pharmaceuticals Corporation; Pfizer Inc.; Piramal Imaging; Servier; Takeda Pharmaceutical Company; and Transition Therapeutics. The Canadian Institutes of Health Research is providing funds to support ADNI clinical sites in Canada. Private sector contributions are facilitated by the Foundation for the National Institutes of Health (www.fnih.org). The grantee organization is the Northern California Institute for Research and Education, and the study is coordinated by the Alzheimer’s Therapeutic Research Institute at the University of Southern California. ADNI data are disseminated by the Laboratory for Neuro Imaging at the University of Southern California.

